# Fascin1 empowers YAP mechanotransduction and promotes cholangiocarcinoma development

**DOI:** 10.1101/2020.11.18.388397

**Authors:** Arianna Pocaterra, Cindy Ament, Silvia Ribback, Xin Chen, Matthias Evert, Diego F. Calvisi, Sirio Dupont

## Abstract

Mechanical forces control cell behavior, including cancer progression. Cells sense forces through actomyosin and YAP, but what regulators of actin mechanotransduction play relevant roles in vivo remains unclear. Here we identify the Fascin1 F-actin bundling protein as a key factor sustaining YAP activation in response to ECM mechanical cues. This is relevant in the mouse liver, where Fascin1 regulates YAP-dependent hepatocyte dedifferentiation. Moreover, Fascin1 is required in the AKT/NICD system and sufficient together with AKT to induce cholangiocarcinomas in mice, recapitulating genetic YAP requirements, and its expression in intrahepatic cholangiocarcinomas correlates with aggressiveness and poor patient prognosis. We propose that Fascin1 represents a pro-oncogenic mechanism that can be exploited during intrahepatic cholangiocarcinoma development to overcome a mechanical tumor-suppressive environment.

## Introduction

Tissue mechanical properties are increasingly considered as key regulators of cell behavior (1,2). Cells are subjected to multiple forces including stretching or compression due to tissue deformation, shearing by liquids flowing at the cell surface, and visco-elastic forces by the extracellular matrix (ECM). Cells can sense these forces and actively respond to them by activating intracellular mechanotransduction pathways, which in turn drive the most appropriate biological response(s): pioneering work indicated how ECM stiffness is sufficient to drive cell behavior, including the choice between proliferation, differentiation or death (3). Moreover, ECM stiffness is also considered an important tissue property that may contribute to the development and progression of cancer (4,5).

ECM mechanotransduction is based on the ability of cells to actively develop actomyosin tension through integrin-mediated adhesions, regulated by RHO GTPases, ROCK (Rho-associated kinases) and MLCK (myosin light chain kinases) (1,6,7). If the ECM opposes strong visco-elastic resisting forces, such as in the case of a stiff ECM, tension across focal adhesions will rise, which will activate a corresponding biological response, often including proliferation and survival. If the ECM instead opposes weak resistance, this will not enable the development of cell tension, leading to cell quiescence and/or to induction of apoptosis. One main mechanism by which ECM forces regulate cell behavior is the regulation of the YAP (yes-associated protein 1) and TAZ (WW-domain transcription regulator 1) proteins (8–10). YAP and TAZ are orthologous transcriptional coactivators that shuttle between the cytoplasm and the nucleus, where they bind with transcription factors of the TEAD family to regulate transcription (11). Among their functions, YAP/TAZ play a prominent role as oncogenic factors in multiple tissues (12). However, what mechanotransduction players are relevant to drive YAP/TAZ activity in tissues and for cancer development, remains poorly defined.

So far, the in vivo physiological relevance for this pathway is based on experimental alteration of ECM stiffness, such as during tissue fibrosis (13–15), and in the MMP14 metalloprotease knockout mice (16). However, while this is in line with the general concept that altered ECM mechanics controls actin dynamics and YAP/TAZ signaling, manipulation of actin dynamics was indirect in these contexts, and entailed the regulation of other parallel and possibly confounding parameters. Moreover, direct modulation of actin dynamics and cell contractility by genetic means provided so far phenotypes that are unrelated to YAP/TAZ (17–19), indicating unexpected specifities. CAPZ is a recent and noteworthy exception, which was identified as regulator of Yorkie - the fly YAP/TAZ homolog - in large-scale screenings (20,21), and later shown to specifically tune the ability of cells to interpret ECM stiffness (22,23). CAPZ knockout enables activation of non-muscle myosin in cells on a soft ECM but also in vivo, in a soft tissue such as the liver, leading to activation of YAP and its hallmark phenotypes, including organ overgrowth and hepatocyte dedifferentiation into atypical ductal cells (24). Thus, CAPZ plays a key role in ECM mechanotransduction, and its inactivation unveiled the importance of YAP mechanotransduction for liver physiology.

CAPZ is the prototypical F-actin barbed-end capping protein, which restrains actin filament elongation at the barbed end concurrently stabilizes actin barbed ends (25,26). Multiple studies indicate that, in migrating cells, CAPZ promotes the formation of branched actin networks that form lamellipodia at the migrating front (27–29). Moreover, CAPZ prevents formation of filopodia by antagonizing Ena/VASP-mediated F-actin elongation and F-actin bundles stabilized by Fascin1 (28,30–32). Collectively, this evidence suggests that CAPZ regulates the balance between two alternative F-actin networks, favoring Arp2/3-dependent branched over bundled F-actin. Such function of CAPZ has also been observed in yeast, suggesting it is not restricted to cell protrusions and cell migration (33).

Here, starting from this “balancing” function of CAPZ, we identify the Fascin1 F-actin bundling protein as a new factor mediating YAP activation in response to ECM mechanical cues, and provide evidence that this represents a pro-oncogenic mechanism that promotes intrahepatic cholangiocarcinoma development.

## Results

### Ena/VASP and Fascin1 sustain YAP/TAZ activity

Evidence for CAPZ regulating the balance between branched and bundled F-actin mainly comes from studies on cell migration, and whether this is relevant for mechanotransduction remains unexplored. To test this, we started our study by testing the functional relevance of proteins promoting the formation of bundled F-actin, such as Formins, Ena/VASP and Fascin1, by using YAP/TAZ as downstream read-out of ECM mechanotransduction. We started our analysis in MCF10A mechano-sensitive mammalian epithelial cells as these are a main cell model for the Hippo pathway and for YAP mechanotransduction (9,22,34–36). The role of formins downstream of integrin/RHO signalling and in mechanotransduction is well known (1), and inhibition of formins with the SMIFH2 small molecule in cells cultured on plastic (i.e. on a stiff substratum) results in nuclear exclusion of YAP/TAZ (Fig. 1A), in line with previous data (22). The role for Ena/VASP proteins is less understood. We thus transfected cells with F4P-GFP-Mito expressing plasmid, sequestering endogenous Ena/VASP proteins at the surface of mitochondria and thus blocking their function (30). As shown in Fig. 1B, expression of F4P-GFP-Mito, but not of its A4P-GFP-Mito control, is sufficient to inhibit cell spreading, as shown by phalloidin staining, and to cause a consequent inhibition of YAP/TAZ, as gauged by nuclear/cytoplasmic localization.

**Figure 1.**
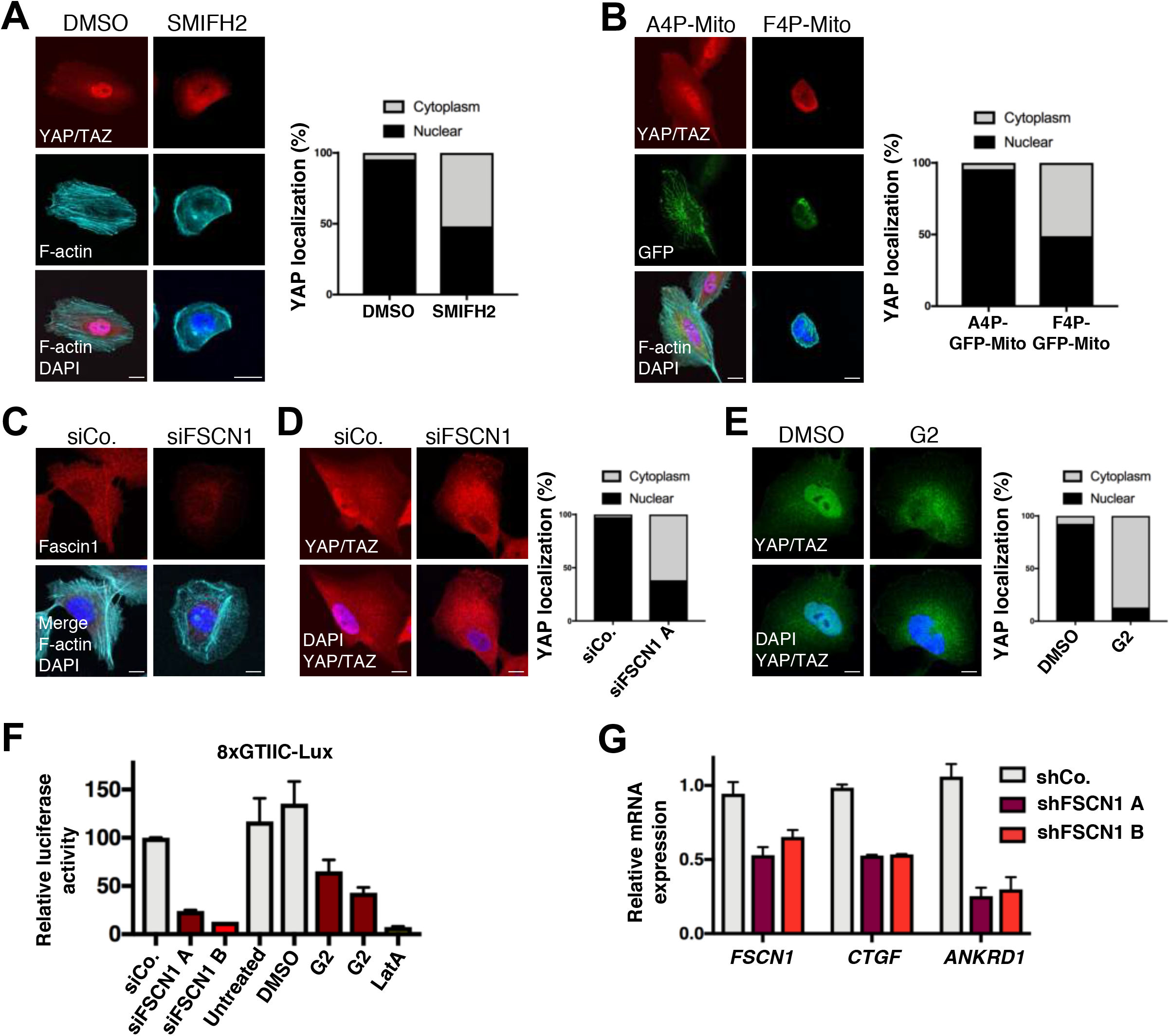
Bundled actin promoted by Formins, Ena/VASP and Fascin1 sustain YAP activity on a stiff substratum. A. Immunofluorescence of MCF10A cells treated with 60μM SMIFH2 Formin inhibitor or with the same amount of vehicle (DMSO) for 24 hours and stained for YAP/TAZ, F-actin (phalloidin) and DAPI as nuclear counterstain. Here and throughout the manuscript, close-up images of single cells representative of the phenotypes are shown, and the quantification of YAP/TAZ subcellular localization is given (on the right), based on multiple pictures of independent single cells, and expressed as percentage of cells in which the nucleus had a stronger staining than the surrounding cytoplasm (Nuclear) or equal/lower than the cytoplasm (Cytoplasm). n=3 (>40 cells per condition in total). *p*=0.013 by unpaired Welch’s t-test. Scale bar= 10μm B. Representative immunofluorescence images of MCF10A cells transfected with a plasmid encoding for the Ena/VASP inhibitor F4P-GFP-Mito, or for its mutated control A4P-GFP-Mito, and stained for YAP/TAZ, F-actin (phalloidin) and DAPI as nuclear counterstain. On the right, quantification of YAP/TAZ subcellular localization as in A. n=3 (>30 cells per condition in total). *p*=0.036 by unpaired Welch’s t-test. Scale bar- 10μm C. Representative immunofluorescence images of MCF10A cells transfected with FSCN1 siRNA (siFSCN1) or control siRNA (siCo.) and stained for Fascin1, F-actin (phalloidin) and DAPI. Scale bar= 10μm D. Representative immunofluorescence images of MCF10A cells transfected with FSCN1 siRNA (siFSCN1) or control siRNA (siCo.) and stained for YAP/TAZ and DAPI, to visualize YAP cellular localization. On the right, quantification of YAP/TAZ subcellular localization as in A. Similar results were obtained with an independent siRNA targeting FSCN1 (not shown). n=3 (>40 cells per condition in total). *p*=0.0283 by unpaired Welch’s t-test. Scale bar= 10μm E. Representative immunofluorescence images of MCF10A cells treated with 100μM G2 Fascin inhibitor (Huang et al., 2015) or its vehicle (DMSO) for 24 hours and stained for YAP and DAPI to visualize YAP cellular localization (quantified on the right as in A). n=3 (>40 cells per condition in total). *p*=0.0088 by unpaired Welch’s t-test. Scale bar= 5μm F. 8XGTIIC-luciferase reporter assay for YAP/TAZ in MDA-MB-231 cells transfected with two independent siRNAs targeting FSCN1 (siFSCN1 A and siFSCN1 B) or control siRNA (siCo.), or treated with different doses of G2 (50μM or 100μM), latrunculinA (LatA 0.5μM) as positive control for inhibition of YAP/TAZ, or DMSO as vehicle. Mean expression in controls was set to 100 and all other samples are relative to this. Data are mean and s.d. G. qPCR for YAP target genes (*CTGF* and *ANKRD1*) in EO771 mouse breast cancer cells stably expressing control short hairpin RNA (shCo.) or two different short-hairpin RNAs targeting FSCN1 (shFSCN1 A and shFSCN1 B). qPCR for *FSCN1* validates efficient downregulation by both short-hairpin RNAs. Data are relative to *GAPDH* expression. Mean expression levels in control cells were set to 1, all other samples are relative to this. Data are mean and s.d. EO771 cells were used to prevalidate shRNAs targeting mouse FSCN1 for subsequent in vivo studies (see below).

Besides Ena/VASP proteins, another key component regulated by CAPZ is Fascin (28). Fascin is a highly conserved protein, encoded by three orthologous genes in mammals, that promotes in vitro the formation of rigid and contractile bundles (37), and is found in filopodia but also around the nucleus (38–42). Fascin1 is the isoform with the wider expression in mice, while Fascin2 and Fascin3 expression is limited to the retina and testis, respectively (43). Recent data suggested a role for Fascin1 as regulator of the Hippo pathway in WM793 melanoma cells and in A549 non-small cell lung cancer cells (44,45), but the functional relevance for this regulation in vivo and in the context of mechanotransduction has not been addressed. MCF10A cells express undetectable levels of *Fascin2* and *Fascin3* mRNA, as measured by qPCR, but express *Fascin1* (Supp. Fig. 1A). We thus knocked-down Fascin1 by RNA interference (Fig. 1C and Supp. Fig. 1B) and found that YAP/TAZ was more cytoplasmic (Fig. 1D). This was independently confirmed by treating cells with the G2 small-molecule inhibitor of Fascin (46) (Fig. 1E). In line, in MDA-MB-231 breast cancer cells that display high level of YAP/TAZ activity (9,47), and whose metastatic ability depends on Fascin (46), inhibition of Fascin1 reduced YAP/TAZ transcriptional activity measured by the established 8XGTIIC-lux luciferase reporter assay (Fig. 1F). Similar results were obtained by monitoring endogenous YAP/TAZ target genes by qPCR in mouse E0771 breast cancer cells stably expressing Fascin1 shRNAs (Fig. 1G), that we used to validate mouse shRNAs to be used in vivo (see below). Collectively, this data indicates that the pool of bundled F-actin promoted by Ena/VASP and Fascin1 sustains YAP/TAZ activity when cells are on a stiff substratum.

### CAPZ and Arp2/3 antagonize Fascin1-dependent actin during mechanotransduction

We and others previously described CAPZ as a YAP/TAZ inhibitor in the context of ECM mechanotransduction, but whether this relates with the ability of CAPZ to promote branched F-actin networks and oppose Fascin1 (see introduction and Supp. Fig. 2A) remains unknown. To test this hypothesis, we inhibited the Arp2/3 complex, the master regulator of branched actin, by treating cells plated on a soft ECM - i.e. the condition in which CAPZ is relevant (22) - with the CK-869 small molecule (48). As shown in Fig. 2A (and quantified in Fig. 2D), CK-869 treatment rescued YAP/TAZ nuclear localization in cells cultured on a soft ECM, and recapitulated the effect of CAPZ depletion (Fig. 2A). Of note, this was specific because CK-869 treatment on plastics, where F-actin bundles are potently sustained by stiffness, did not change YAP/TAZ localization (Supp. Fig. 2A and B), similar to previous observations on CAPZ (22). Overall, this data indicates that CAPZ and Arp2/3, regulators of branched F-actin, contribute in keeping YAP/TAZ out of the nucleus on a soft ECM.

**Figure 2.**
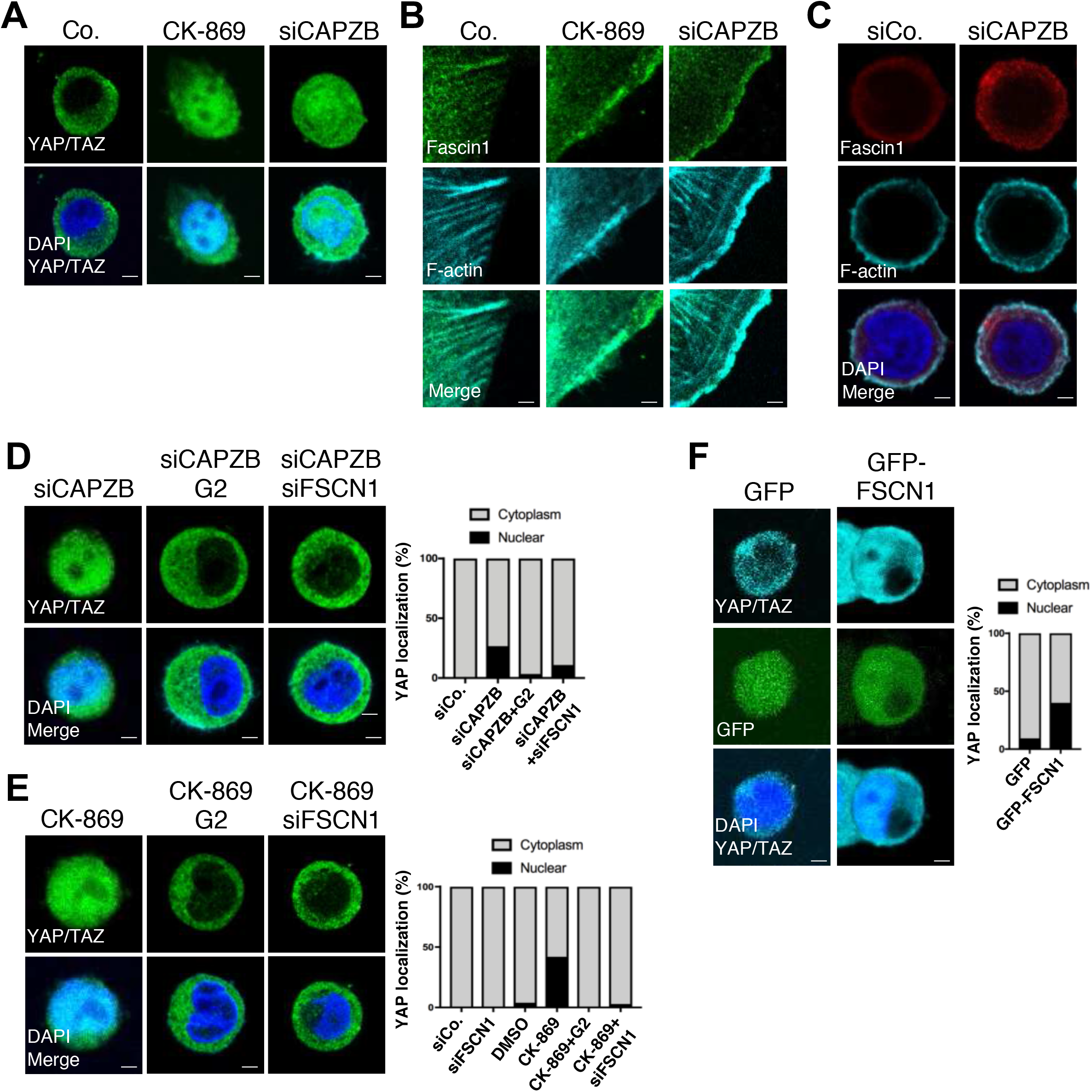
CAPZB and Arp2/3 inhibition induce nuclear YAP on soft ECM substrata, which depends on Fascin1. A. Representative immunofluorescence images of MCF10A cells plated on soft Fibronectin-coated polyacrylamide hydrogels (E≈0.5kPa) for one day and then treated for additional 24 hours with 80μM CK-869 (an established Arp2/3 inhibitor) or vehicle control (DMSO). Where indicated cells were transfected with CAPZB siRNA (siCAPZB). Cells were stained for YAP/TAZ and DAPI as nuclear counterstain. See quantification of YAP/TAZ subcellular localization in D. Scale bar= 3μm B. Representative immunofluorescence images of MCF10A cells treated with the CK-869 Arp2/3 inhibitor or vehicle control (DMSO) or transfected with CAPZB siRNA (siCAPZB). Cells were stained for Fascin1 and F-actin (phalloidin). Scale bar= 2μm C. Representative immunofluorescence images of MCF10A transfected with CAPZB siRNA (siCAPZB) or control siRNA (siCo.) and replated on soft hydrogels for 48 hours, stained for Fascin1 and F-actin (phalloidin). Scale bar= 2μm D and E. Representative immunofluorescence images of MCF10A cells transfected with CAPZB siRNA (siCAPZB, D) or treated with 80μM CK-869 (an established Arp2/3 inhibitor, E) and cultured on soft hydrogels. Where indicated cells were co-treated with the G2 Fascin inhibitor or co-transfected with Fascin1 siRNA (siFSCN1). Cells were stained for YAP/TAZ and DAPI. On the right, quantification of YAP/TAZ subcellular localization for the indicated conditions. n=3 (>25 cells per condition in total). Scale bar= 2μm F. Representative immunofluorescence images of MCF10A cells transfected with FSCN1 and GFP (GFP-FSCN1) or GFP plasmid alone, and cultured on soft hydrogels for 24 hours. On the right, quantification of YAP/TAZ subcellular localization. n=3 (>20 cells per condition in total). p=0.016 by unpaired Welch’s test. Scale bar= 4μm

Given the potential competition between branched and bundled F-actin networks (49,50), we then tested whether Arp2/3 and CAPZ regulate YAP/TAZ by regulating Fascin1. In line with this hypothesis, treatment of cells with CK-869 or depletion of CAPZ led to accumulation of Fascin1-positive structures at cell edges (Fig. 2B and Supp. Fig. 2B), where CAPZ is mainly localized (Supp. Fig. 2C). These structures in part co-localized with Vinculin, a marker for mature focal-adhesions (Supp. Fig. 2D). Moreover, enhanced Fascin1 staining was observed in CAPZ-depleted cells plated on a soft ECM (Fig. 2C). Most importantly, Fascin1 activity is required for nuclear YAP/TAZ induced by Arp2/3 inhibition or by CAPZ knockdown (Fig. 2D and E). Collectively, these results suggest that when cells are on a soft ECM, CAPZ and Arp2/3 oppose YAP/TAZ nuclear localization, and that this occurs because they inhibit Fascin1 and the corresponding bundled F-actin (see Suppl. Fig. 2A). To reinforce this idea, overexpressing active Fascin1 in cells on a soft ECM was sufficient to promote partial YAP/TAZ nuclear accumulation (Fig. 2F).

### CAPZ antagonizes Fascin1-dependent YAP activity in the liver

To challenge the relevance of our findings in vivo, we sought to transpose our observations to the mouse liver, a model system to study YAP/TAZ biology (51,52), and an organ where cells lay in a soft environment (53,54). Moreover, in this tissue, the hepatocyte-specific inactivation of *Capzb* leads to activation of YAP mechanotransduction, proliferation and dedifferentiation of hepatocytes into atypical ductal cells (ADC) of cholangiocellular identity (24).

We initially tested whether Fascin1 is downstream of CAPZ also in vivo. For this we inhibited Fascin1 in liver cells by administering the G2 Fascin inhibitor by i.p. injection to CAPZ LKO mice, and scored hepatocyte dedifferentiation as read-out of YAP function (24,52,55). As shown in Fig. 3A, G2 treatment restricted the expansion of the ADC marker CK19 in CAPZ LKOs. The effect was partial, likely due to the relatively low affinity of G2 for Fascin1 (46). To test whether Fascin1 was sufficient to activate YAP in vivo, we then overexpressed Fascin1 in the liver of adult wild type mice by using hydrodynamic tail vein (HTV) injection of transposon plasmids and scored established YAP-induced phenotypes. As shown in Fig. 3B, Fascin1 was sufficient to increase proliferation, as shown by EdU incorporation, and to induce ADCs, as gauged by staining for the A6 cholangiocellular marker. Importantly, these phenotypes were prevented when *Yap1/Wwtr1(TAZ)* were knocked-out in Fascin1-expressing hepatocytes (Fig. 3C), indicating an effect mediated by cell-autonomous activation of YAP/TAZ. These results suggest that CAPZ maintains hepatocyte cell fate by inhibiting Fascin1-dependent YAP activation.

**Figure 3.**
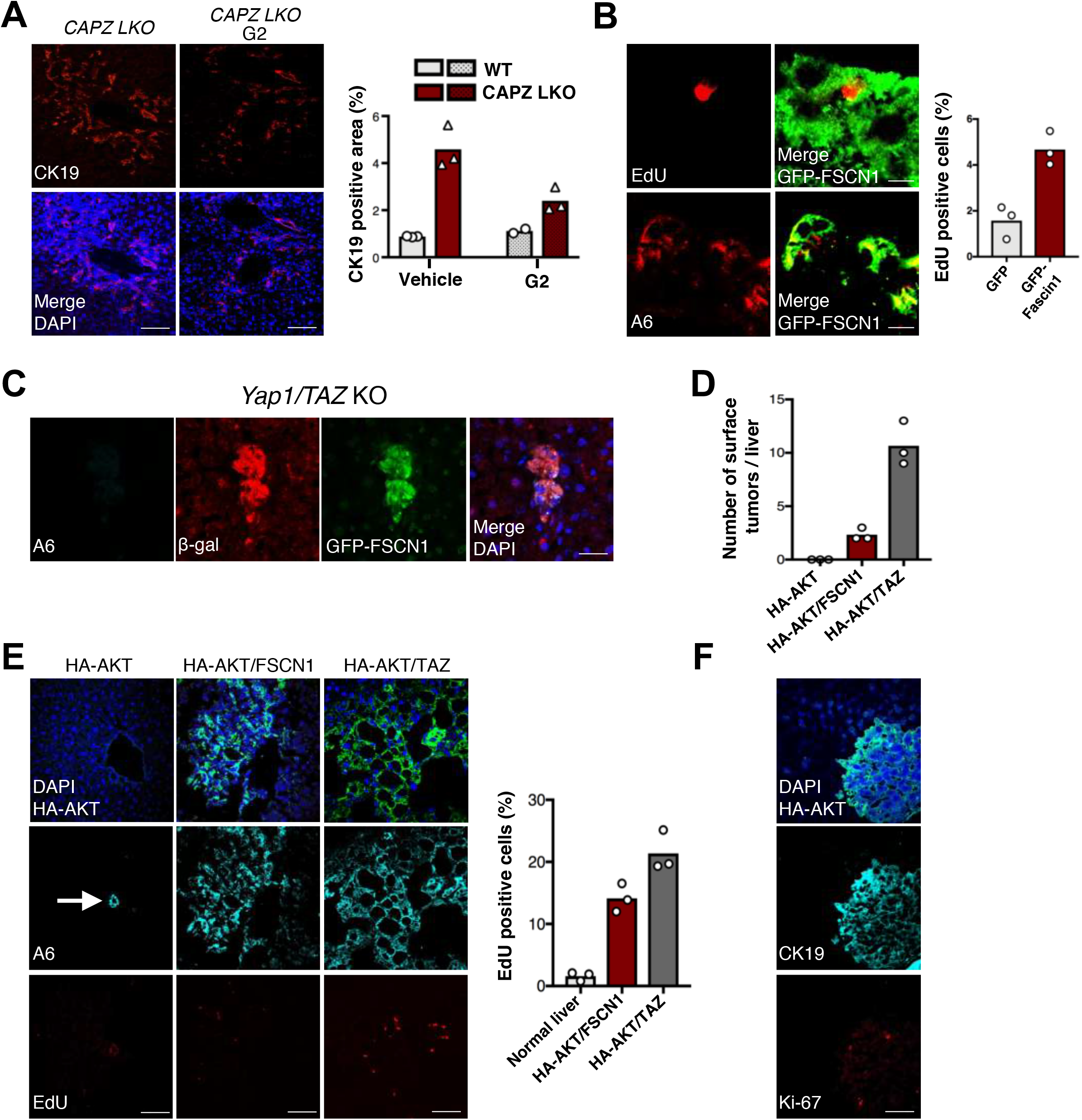
Fascin1 regulates hepatocyte cell fate through YAP/TAZ and promotes cholangiocarcinoma development. A. Representative immunofluorescence stainings of liver sections from adult tamoxifen-injected *Albumin-CreERT2; Capzbfl/fl* mice (*CAPZ LKO*) mice injected i.p. with the G2 Fascin inhibitor or with vehicle (5% DMSO), stained for the cholangiocellular marker CK19 and DAPI as nuclear counterstain. Dedifferentiation of hepatocytes into ADC in *CAPZ LKO* depends on *Yap1* and on increased actomyosin contractility (Pocaterra et al., 2019). Analyses were carried out 15 days after tamoxifen injection. Induction of CK19-positive cells in CAPZ LKO was confirmed compared to non-injected mice (not shown). On the right, quantification of CK19-positive area in sections of the portal area for the indicated conditions. Average and single data (mice, n=3). KO vs. KO+G2 *p*=0.033 by unpaired Welch’s test. Scale bar= 100μm. B. Representative immunofluorescence stainings of liver sections from C57BL/6N mice transduced by hydrodynamic tail vein (HTV) injection with transposon plasmids encoding for GFP-Fascin1. Liver sections were stained for EdU (to visualize hepatocytes in S-phase) and the cholangiocellular marker A6. Analyses were carried out 15 days after HTV injection. On the right, quantification of EdU-positive hepatocytes in livers transduced with GFP-Fascin1 or with GFP alone, which served as experimental control (pictures not shown). Mean and single points (mice, n=3). *p*=0.049 by unpaired Welch’s test. Scale bar = 5μm. C. Representative immunofluorescent stainings of liver sections from *Yap1fl/fl; Wwtr1(TAZ)fl/fl; ROSA26-LSL-lacZ* mice transduced by HTV with transposon plasmids encoding for GFP-Fascin1 and CRE recombinase (to induce *Yap1/TAZ* knockout), stained for the cholangiocellular marker A6, β-galactosidase (β-gal - lineage tracer for CRE-induced recombination), GFP and DAPI as nuclear counterstain. Analyses were carried out 15 days after HTV injection and induction of CRE activity by tamoxifen. *Yap1fl/fl; Wwtr1(TAZ)fl/fl; ROSA26-LSL-lacZ* mice without CRE recombinase are wild-type; injection of GFP-Fascin1 in *Yap1fl/fl; Wwtr1(TAZ)fl/fl; ROSA26-LSL-lacZ* mice induced the same phenotypes observed in B (not shown). Scale bar= 10μm. D. Number of macroscopic tumor lesions detected at the liver surface in C57BL/6N mice transduced by HTV injection with transposon plasmids expressing myristoylated-HA-AKT alone, together with Fascin1 (AKT/FSCN1), or together with activated TAZ (AKT/TAZ). Livers were analyzed 7 months after HTV. Mean and single points (mice, n=3). AKT vs. AKT/FSCN1 *p*=0.019, AKT vs. AKT/TAZ *p*=0.013, AKT/FSCN1 vs. AKT/TAZ *p*=0.015, by unpaired Welch’s tests. E. Representative immunofluorescence images of liver sections from C57BL/6N mice transduced by HTV injection as in D. Liver sections were stained for HA to localize AKT-expressing neoplastic lesions, EdU to visualize cells in S-phase, the cholangiocellular marker A6, and DAPI as nuclear counterstain. The white arrow on the left panel indicates a normal A6-positive bile duct. Elimination of cells expressing only AKT is likely due to the higher resistance to oncogenic transformation of the C57BL/6N strain compared to FVB/N. Livers were analyzed 7 months after HTV. On the right: quantification of EdU-positive cells in liver sections. Average and single points (mice, n=3). WT vs. FSCN1 *p*=0.0067, WT vs. TAZ *p*=0.0069, FSCN1 vs. TAZ *p*=0.040, by unpaired Welch’s tests. Scale bar= 80μm. F. Representative immunofluorescence images of liver sections from AKT/FSCN1 mice. Liver sections were stained for HA to localize neoplastic lesions, Ki67 to visualize proliferating cells, the cholangiocellular marker CK19 and DAPI as nuclear counterstain. Scale bar= 60μm.

### Fascin1 has a pro-oncogenic function in cholangiocarcinomas

In the liver, activation of YAP not only induces transdifferentiation of hepatocytes into ADCs, but also promotes the development of hepatocellular carcinomas (HCC) and intrahepatic cholangiocarcinomas (iCCA) (56–59). The effects of Fascin1 on YAP activity and ADC formation shown above prompted us to explore the role of Fascin1 for liver tumorigenesis. Expression of Fascin1 together with myristoylated AKT (myrAKT), a known driver of liver tumors (60), was sufficient to promote the formation of macroscopic masses, while cells expressing only myrAKT were lost in the same time-frame, likely due to insufficient fitness in the C57BL/6N background (Fig. 3D). myrAKT/FSCN1 tumors display enhanced proliferation as gauged by EdU incorporation, and were of cholangiocellular identity because they express the A6 and CK19 markers (Fig. 3E and F). Of note, a similar phenotype can be also obtained by expression of myrAKT together with activated YAP (61) or TAZ (Fig. 3E). Thus, Fascin1 cooperates with activated AKT to induce liver neoplasia. Moreover, as we failed to observe myrAKT/Fascin1-positive and A6- or CK19-negative lesions (not shown), this suggests that Fascin1 skews the ability of myrAKT to induce both HCC and iCCA (62) specifically towards the iCCA subtype.

To explore the functional requirement of Fascin1 for iCCA, we next explored whether experimental iCCA induced in mice by different oncogene combinations display increased Fascin1 expression. For this we stained livers of FVB/N mice with iCCA induced by hydrodynamic tail vein (HTV) injection of myrAKT together with activated N-Ras-V12D (myrAKT/N-Ras), with activated Notch Intracellular Domain (myrAKT/NICD) or with activated YAP-S127A (myrAKT/YAP) (55,60,61,63,64). In all models tested, a strong Fascin1 immunoreactivity was detected in endothelial cells, and used as an internal control of the staining. However, only the myrAKT/NICD combination displayed consistent, strong and diffuse cytoplasmic Fascin1 expression in tumor cells when compared to the surrounding normal tissue (Fig. 4A, see higher magnifications in Supp. Fig. 3A). Absence of Fascin1 overexpression in other oncogenic combinations likely reflect different underlying molecular mechanisms. Tumors induced by NICD alone were also negative for Fascin1 expression in tumor cells, suggesting the need for combined NICD and AKT signaling (Fig. 4A). Of note, development of iCCA by myrAKT/NICD relies on endogenous YAP in cancer cells (55,58), thus representing the ideal experimental set-up to test the requirement for Fascin1. We thus combined myrAKT/NICD expression together with control shRNA (shCo.) or with a pre-validated Fascin1 shRNA (shFascin1 - see above). myrAKT/NICD+shCo. caused the appearance of multiple focal neoplasms 3 weeks after injection (Fig. 4B and Supp. Fig. 3B), in line with previous evidence (Fan et al., 2012), and Fascin1 shRNA caused a significant reduction in the number and size of lesions (Fig. 4B and C). This data indicates that Fascin1 is required in the myrAKT/NICD system, and sufficient together with myrAKT, for the development of intrahepatic cholangiocarcinomas in mice.

**Figure 4.**
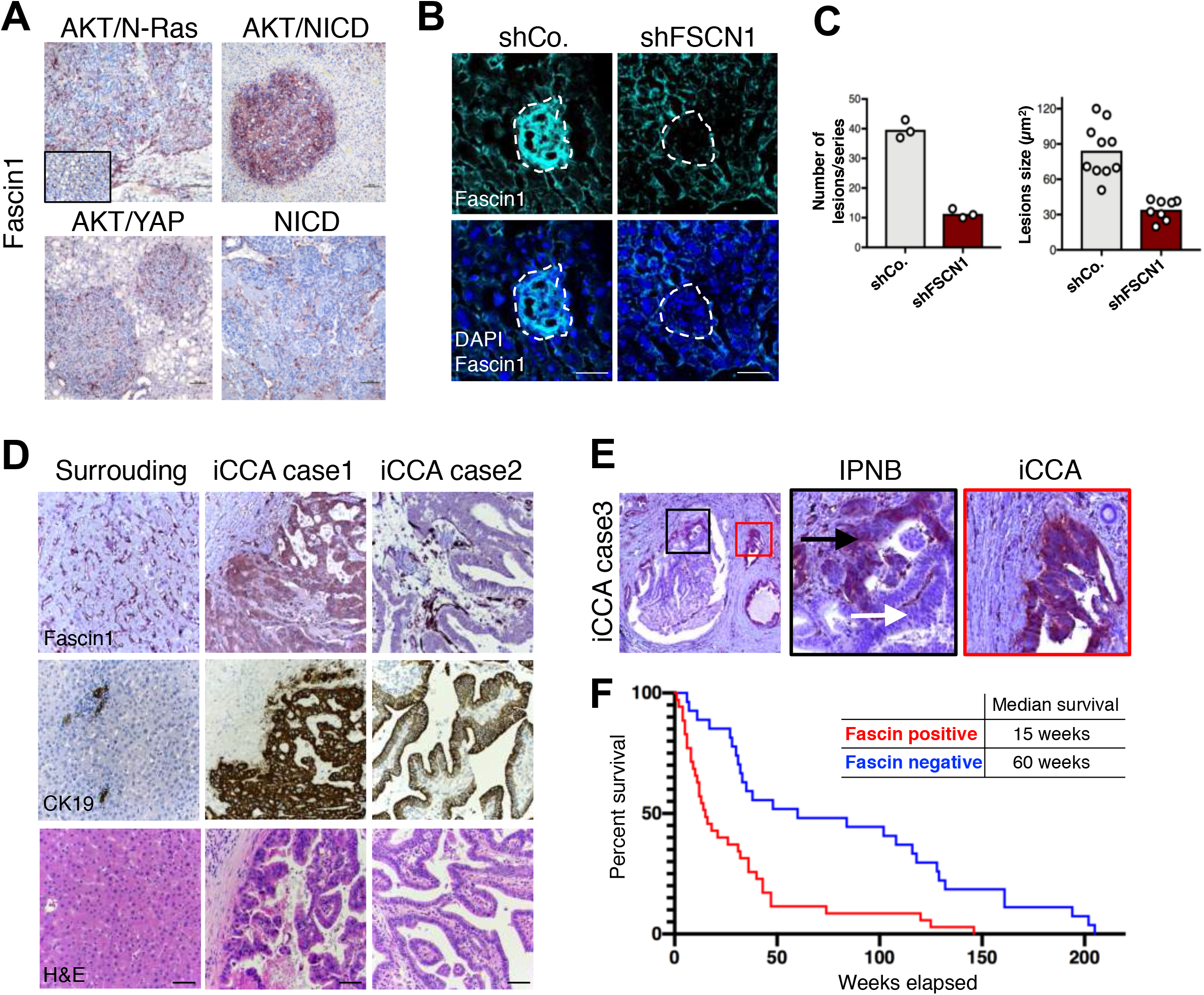
Fascin1 is overexpressed in intrahepatic cholangiocarcinomas and required for disease progression. A. Representative Fascin1 immunohistochemistry in iCCA formed in FVB/N mice transduced by hydrodynamic tail vein (HTV) injection with transposon plasmids encoding for Notch Intracellular Domain (NICD), or myristoylated-AKT together with NICD (AKT/NICD), with N-Ras V12D (AKT/N-Ras), and with YAP S127A (AKT/YAP). Fascin1 immunoreactivity was found to be limited to liver sinusoids, stromal, and endothelial cells in the normal tissue and in most iCCA models, including the hepatocellular carcinomas developed in the AKT/N-Ras mice (inset). In contrast, cholangiocellular lesions developing in AKT/NICD mice exhibited intense cytoplasmic staining for Fascin1 in tumor cells. n=5 mice for each model were consistent. Original magnifications: 100x; scale bar= 100μm. B. Representative immunofluorescence images of liver sections from C57BL/6N mice transduced by HTV injection with HA-AKT/NICD and control short-hairpin RNA (shCo.) or with a short-hairpin RNA targeting Fascin1 (shFSCN1). Liver sections were stained for HA (see Supp. Fig. 4B), endogenous Fascin1 and DAPI as nuclear counterstain. Livers were analyzed 3 weeks after HTV. Note how this validates the antibody used in D and E below. Scale bar= 60μm. C. Quantifications of the number and size of liver lesions shown in B. On the left: mean and single points (mice, n=3) of the total number of HA-positive lesions in 5 non-consecutive sections from each liver (series), *p*=0.0008 by unpaired Welch’s t-test. On the right: mean and single points of HA-positive lesions’ size from n=3 mice, *p*<0.0001 by Mann-Whitney test. D. Representative Fascin1 immunohistochemistry on human iCCA. In the non-neoplastic liver (surrounding liver), Fascin1 immunoreactivity is limited to liver sinusoids and endothelial cells. iCCA case 1 shows intense cytoplasmic staining for Fascin1 in tumor cells, and is representative of 35/62 (56%) cases. iCCA case 2 displays immunolabeling for Fascin1 in vascular structures but not in tumor cells. CK19 immunoreactivity was used to demonstrate the cholangiocellular differentiation of the iCCA lesions. Original magnification: 200x; scale bar=100μm. E. Left panel: low magnification of a liver section stained for Fascin1, containing a malignant pre-invasive condition (black inset, intraductal papillary neoplasm of the bile duct, IPNB) and a frankly invasive iCCA lesion (red inset) adjacent to each other. Black panel: higher magnification of the IPNB, showing immunoreactivity for Fascin1 limited to the invasion front (black arrow), but otherwise negative (white arrow). 4/13 cases were positive. Red panel: higher magnification of the iCCA, exhibiting strong and homogeneous staining for Fascin1 in the cytoplasm of tumor cells. 9/13 cases were positive. Original magnification: 200x; scale bar= 100μm. F. Kaplan-Meier survival analysis of human iCCA patients based on Fascin1 expression levels. Tumors positive for Fascin1 staining have poorer outcomes (median survival 15 weeks) compared with negative ones (median survival 60 weeks). n=62. *p*=0.0001 by log rank Mantel-Cox test.

Finally, we sought to understand whether these data bear significance for human liver disease. We performed Fascin1 immunohistochemistry with our extensively validated antibody (see Fig. 1C and 4B) on a series of human liver cancers with matched normal liver tissues (Supp. Tables 1 and 2). In the normal human liver Fascin1 staining was restricted to vascular structures, with negative or very low staining in hepatocytes (Fig. 4D), as also seen in mice; we instead observed strong diffuse cytoplasmic staining in 35/62 (56%) of human iCCA (Fig. 2K), suggesting the overexpression observed in mouse myrAKT/FSCN1 tumors finds a correlate in the human disease. We also stained human HCC, and in this case only 13/50 (26%) were positive (Supp. Fig. 3C). We then took advantage of a collection of human iCCA cases with adjoining preinvasive and invasive lesions, were we noted less-frequent (4/13) and restricted expression of Fascin1 to cells at the invasive front of preinvasive neoplasms, which became more frequent (9/13) and homogeneous in invasive carcinomas (Fig. 4E). All cancer cells were positive for YAP immunostaining suggesting that in human iCCA, like in mouse experimental iCCA (see references above), multiple different mechanisms converge on sustaining YAP levels. Finally, to explore a correlation between Fascin1 expression and prognosis, we estimated the survival curves of patients whose tumor cells stained positive or negative for Fascin1 by the Kaplan-Meier method, and found a striking correlation with poor prognosis in iCCA (Fig. 4F, see also Table 1 and 2), with an estimated median survival of 60 weeks for Fascin1-negative patients and of 15 weeks for Fascin1-positive patients. Together with data in mice, this suggests that Fascin1 expression this suggests that Fascin1 expression is enhanced in a proportion of human iCCA, and that it represents a significant factor that promotes disease progression.

## Discussion

The identification of the YAP mechanotransduction system provided a powerful model to study the effects of tissue mechanical properties on cell behavior. Yet, despite a wealth of knowledge on what are the most upstream players of this pathway (i.e. at the level of focal adhesions) (1,6,65) and some very recent hints on what are the mechanisms proximal to YAP regulation (66–69), what are the relevant intermediate players, and what actin structures are involved, remains largely unknown. Moreover, which of these players are important to drive YAP mechanotransduction in vivo, and whether they may play an active role in cancer progression, remains an even less charted territory.

Here, starting from known but so far YAP- and mechano-unrelated functions of CAPZ in shifting the balance between actin bundled vs. branched structures during cell migration (28,30), we identify Ena/VASP and Fascin1 proteins as required for YAP activity in response to ECM stiffness. Conversely, on a soft ECM CAPZ and Arp2/3 complexes, promoters of branched F-actin, inhibit YAP activity by limiting the formation of Fascin1-dependent actin. In line, expression of activated Fascin1 is sufficient to drive nuclear YAP on a soft ECM. The concept of competition between different actin networks in cells has been previously proposed in yeast, where Profilin regulate the balance between Arp2/3 and Formin activity (50,70), in epithelial cells of liver origin, where actomyosin bundles prevents the formation of Arp2/3 dependent submembranous actin (49), and in axon growth cones, where Arp2/3 activity can restrict myosin-mediated contractility (71). Our data suggests that such competition may also occurs during ECM mechanotransduction, and becomes relevant when cells are in conditions of decreased ECM stiffness: in these conditions, not only RHO signalling is decreased, leading to decreased activity of ROCK/MLCK and of Formins (1), but at the same time CAPZ-dependent and Arp2/3-dependent actin outbalances actin bundles promoted by Ena/VASP and Fascin1.

The role of Ena/VASP and Fascin1, key factors driving formation of filopodia, may indicate a specific function of filopodia in our phenotypes (38,42). Filopodia are mechano-sensitive structures, they support the formation of nascent adhesions that can subsequently develop into mature focal adhesions in response to ECM forces, and they provide migrating cells the ability to probe the mechanics of the microenvironment (1,39,72). However, we did not observe major induction of filopodia upon Fascin1 overexpression, Arp2/3 inhibition or CAPZ inactivation, at least under our imaging resolution, and Fascin1 staining in cholangiocarcinomas appear cytoplasmic diffuse, not limited to the cell periphery/protrusions. Thus, these players might regulate other cytoplasmic bundled actin structures relevant for mechanotransduction.

Importantly, this new activity of Fascin1 is relevant in the liver, a model system to study YAP and mechanotransduction. We found that Fascin1 activation is sufficient to induce mouse hepatocyte dedifferentiation into atypical ductal cells (ADC, also called oval cells or bipotent liver progenitor cells) in a YAP/TAZ-dependent manner, and that Fascin1 is required for ADC formation downstream of CAPZ inactivation, where enhanced contractility drives YAP activation (24). This activity of Fascin1 is coherent with previous findings, which however implied an actin-independent function of actin, and without in vivo relevance (44,45). These findings led us to explore the role of Fascin1 in liver cancer. We found that human intrahepatic cholangiocarcinomas (iCCA) overexpress Fascin1, which correlates with more invasive iCCAs and poor prognosis. Moreover, we found a similar overexpression in iCCAs induced by experimental activation of AKT and Notch in mice, further validating this system as a relevant model for human iCCA, and providing candidate molecular mechanisms mediating Fascin1 overexpression. Regulation of Fascin1 expression observed in tumors appears different from regulation of Fascin1 localization that we observe upon inhibition of CAPZ and Arp2/3, but still sufficient to activate YAP and the corresponding biological responses, as we found that increased Fascin1 is required for the growth of iCCA induced by AKT and Notch, and cooperates with AKT to induce iCCA formation.

We thus propose that Fascin1 represents a novel player of the YAP mechanotransduction machinery whose expression can be selected by some oncogenic lesions to overcome a mechanical tumor-suppressive environment. This might be relevant in the pancreas, where stromal stiffening is an important driver of tumor development (73,74), and where Fascin1 is also genetically required (75). Fascin-specific inhibitors have been developed that can be used in vitro, although the high working concentration of these compounds precludes their efficient use in vivo (46,76,77); the observation that one of such inhibitors dose-dependently blocks the growth of multiple Fascin1-positive human iCCA cell lines in vitro (Supp. Fig. 4) suggests that the development of more active and specific compounds is desirable.

## Methods

### Cell lines, transfections and microfabrications

Human mammary epithelial MCF10A cells (ATCC) were cultured in DMEM/F12 supplemented with 5% Horse Serum, 2mM Glutamine, insulin (Sigma), cholera toxin (Sigma), hEGF (Peprotech) and hydrocortisone (Sigma) as in (Debnath et al., 2003). Human breast cancer MDA-MB-231 cells (ATCC) in DMEM/F12 with 10% FBS and 2mM Glutamine. All cell lines were routinely tested with universal mycoplasma ATCC detection kit 30-1012K and were always negative. Murine breast cancer E0771 cells were cultured in RPMI medium supplemented with 10% FBS, 2mM Glutamine and 1% HEPES. siRNA transfections were done with Lipofectamine RNAi MAX (Invitrogen) and plasmid DNA transfections were done with Transit-LT1 (MirusBio) according to the manufacturer’s instructions. Where indicated, transfections were carried out on plastic vessels and cells were subsequently replated on hydrogels. Sequences of siRNAs are provided in Table 1. Fibronectin-coated polyacrylamide hydrogels (E ≈ 0.5 kPa) were assembled in-house as in Ref.(24).

### Plasmid and reagents

The small-molecule inhibitors were SMIFH2 (Sigma s4826, 60 microM), CK-869 (Sigma c9124, 80 microM) LatrunculinA (Sigma L5163, 0.5 microM), G2 (Xcessbio M60269 for cell treatments, Valuetech custom synthesis for mouse injections). All plasmids were sequence-verified before use and transfected as endotoxin-free maxi preps. siRNAs were selected among FlexiTube GeneSolution 4 siRNA sets (Qiagen) and reordered after validation as dTdT-overhanging 19 nt RNA duplexes (Thermo). shRNAs were selected among pre-validated Mission pLKO1-shRNA (Sigma) and the corresponding U6-shRNA-cPPT cassettes were subcloned into PB-empty vector (24). siRNA and shRNA sequences are as follows:

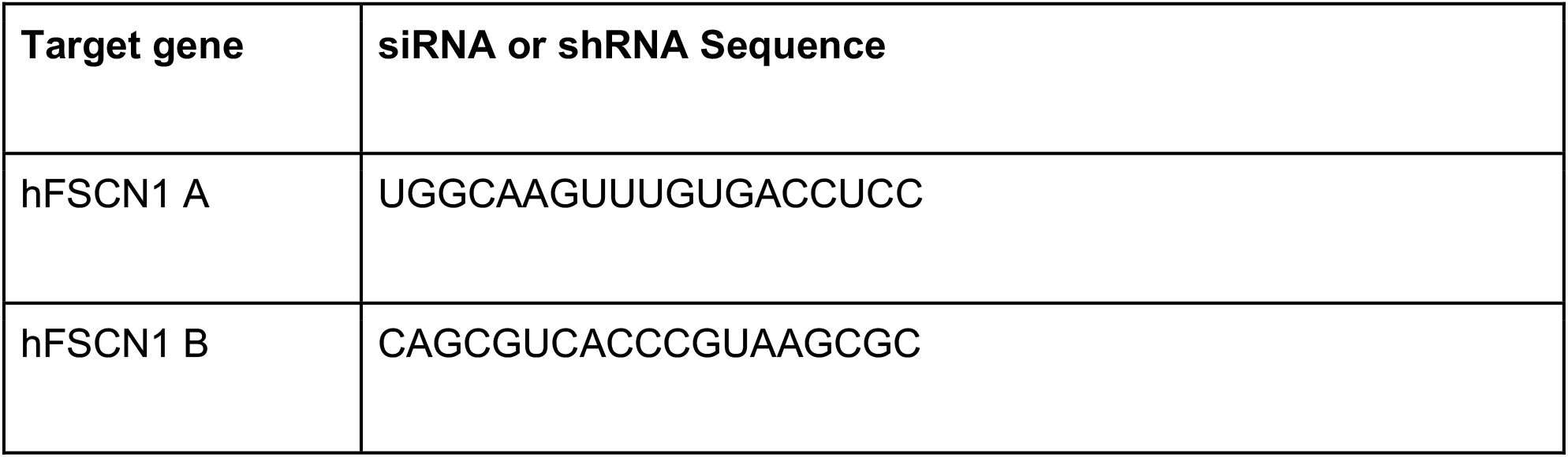
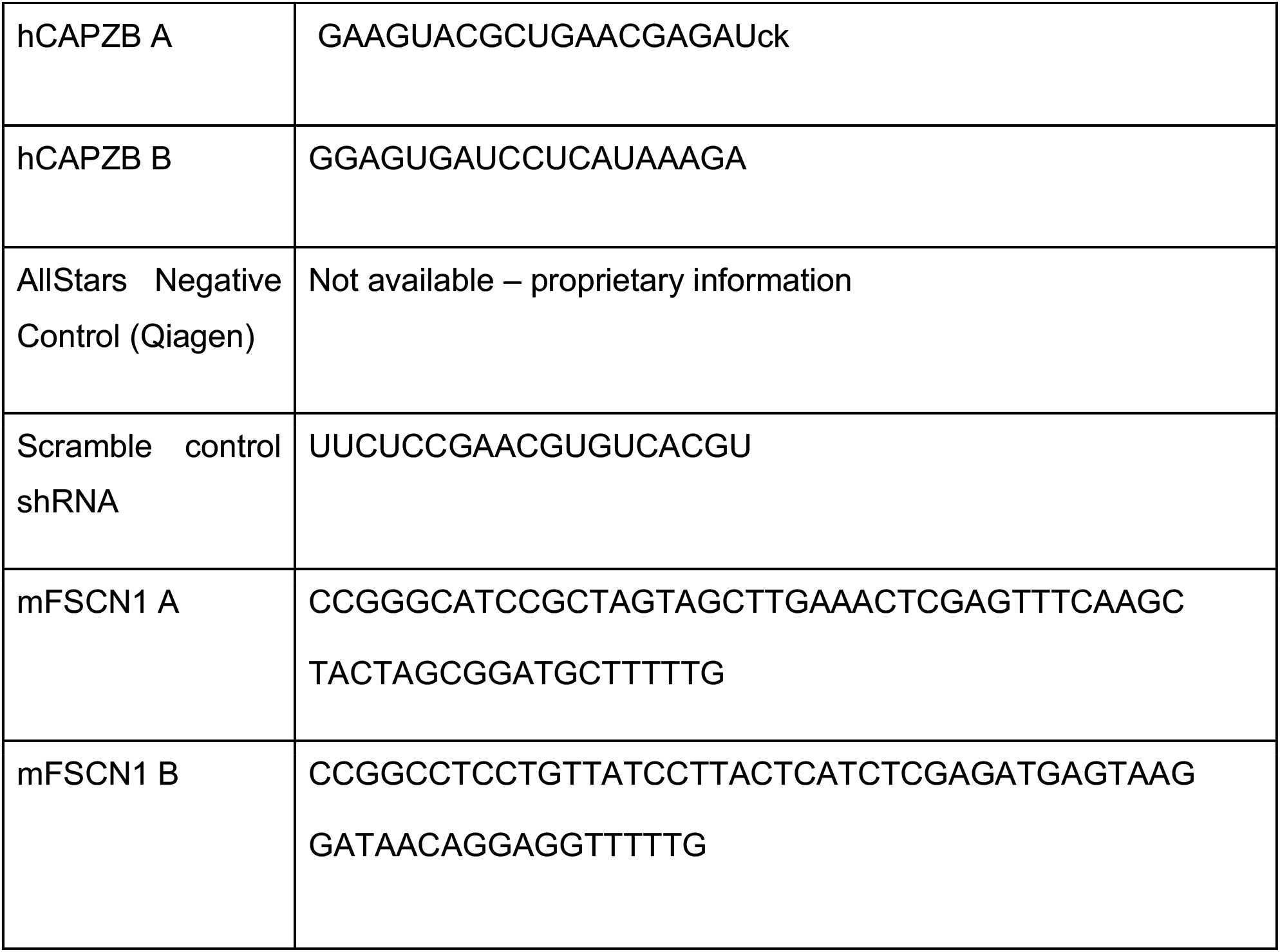

### Human tissue samples

Sixty-two intrahepatic cholangiocarcinomas (iCCA) and fifty hepatocellular carcinomas (HCC) as well as the corresponding surrounding non-tumorous liver tissues were used for the study. Patients’ clinicopathological features are summarized in Table 1 and 2. Tumors were divided in iCCA/HCC with shorter/poorer (iCCA, n = 35; HCC, n = 28) and longer/better (iCCA, n = 27; HCC, n = 28) survival, characterized by <3 and ≥3 years’ survival following partial liver resection, respectively. Liver tissues were collected at the Universities of Greifswald (Greifswald, Germany) and Regensburg (Regensburg, Germany). Institutional Review Board approval was obtained at the local Ethical Committees of the Medical Universities of Greifswald (approval code: BB 67/10) and Regensburg (17-1015-101), in compliance with the Helsinki Declaration. Written informed consent was obtained from all individuals.

### Mice and treatments

Mice were C57BL/6N as in Ref. (24). Sex allocation was random. For G2 treatment, mice were administered 100 mg/kg of G2 by daily i.p. injection, 5 days per week, for 2 weeks, starting together with tamoxifen injection (to induce CAPZB recombination). Animal experiments were performed according to our institutional guidelines as approved by the University Animal Welfare Commission (OPBA) and authorized by the Ministry of Health (945/2015-PR and 54/2015-PR). Reporting was according to the ARRIVE guidelines.

Hydrodynamic tail-vein injection was used to transduce hepatocytes of of 4/6-week-old mice with exogenous DNA. 50μg of total PiggyBac (PB) and/or Sleeping Beauty (SB)-transposon plasmid DNA together with 10μg of hyperactive PB Transposase plasmid DNA (hyPBase) or hyperactive Sleeping Beauty transposase (pCMVT7-SB100, Addgene 34879) were diluted in sterile Ringer’s solution in a volume corresponding to 10% of the body weight (on average, 18-22 grams), and injected via the tail vein in a maximum time of 8-10 seconds. PB-transposon plasmids were obtained by subcloning the cDNAs of interest (GFP, GFP-IRES-Fascin1 S39A, FLAG-mTAZ 4SA) in a PB-empty plasmid.

Mice were euthanized and abdominal contents exposed. Trans-cardiac perfusion (29-gauge needle) with cold 1XPBS (10-20ml) was performed to reduce blood contaminants. The liver was removed and placed in a clean petri dish with 1XPBS on ice. The liver was immediately divided in parts and snap-frozen in liquid nitrogen for extraction of mRNA/proteins or embedded in OCT and stored at −80° for subsequent analyses.

### Antibodies and immunofluorescence

| **Epitope** | **Brand and catalog** | **Validation** |
| --- | --- | --- |
| CK19 | DSHB TROMAIII | Positive IF staining on bile ducts.<br>NOTE: the antibody requires inclusion of liver tissues in OCT without prior fixation with PFA. |
| A6 | DSHB A6 BCM | Positive IF staining on bile ducts. |
| YAP | SC-101199<br>used in vitro | Detects a prominent band in WB of the expected size, which disappears in human cells treated with YAP siRNA.<br>Detects nuclear/cytoplasmic shuttling of YAP in multiple conditions. |
| YAP | PROTEINTECH 13584-1-AP used in vivo | Detects a prominent band in WB of the expected size, which disappears in human cells treated with YAP siRNA. Detects nuclear/cytoplasmic shuttling of YAP in multiple conditions in vitro. IF signal is lost in *YAP*-null liver tissue. |
| FSCN1 (55K-2) | SC-21743 | IF staining shows a specific localization in filopodia and in cytoplasm colocalized with phalloidin. The staining disappears in human cells and mouse tumors expressing FSCN1 siRNAs or shRNAs. |
| FSCN1 | ab126772 | Staining is consistent with the other antibody used in the study |
| HA | SC-7392 | Positive WB or IF staining only in cells transduced with plasmids encoding for HA-tagged proteins |
| β-gal | ab9361 | Positive IF staining only in tissues or cells expressing a lacZ transgene. |

For immunofluorescence, cells were fixed in 4% PFA for 10 minutes. After washes, cells were permeabilized with PBS-Triton 0.5% for 20 minutes following by blocking buffer (PBS-Triton 0.1%, 2% goat serum) incubation for 1 hour. Primary antibody was incubated overnight at +4°. The day after, cells were incubated with Alexa fluor(Invitrogen) - conjugated secondary antibody for 1 hour at room temperature. Dishes were mounted with ProLong™ Gold Antifade Mountant with DAPI (P36935 Thermofisher). Images were acquired with a Leica SP5 or with a ZEISS LSM700 confocal microscope equipped with CCD camera, using Leica LAS AF or ZEN 2 software, or with a standard Leica DM5000B microscope. Raw images were analyzed with ImageJ (Fiji).

For immunofluorescence on liver sections, OCT-embedded tissue was cut into 5-8μm thick sections with a Leica CM1950 cryostat. Sections were dried at RT for 30 minutes on a glass coverslip (VWR), and either stored dried at −80°C or directly processed by rehydration in 1XPBS followed by fixation in 4% PFA for 15 minutes. Permeabilization was performed in 1XPBS-Triton 1% for 20 minutes. Blocking was 10% goat serum in 1XPBS-Triton 0.5% for 1 hour at RT.

For EdU labelling, mice were injected with 12.5mg/kg of EdU in sterile 1XPBS (A10044 Molecular Probes) 15 hours before tissue sampling. Cells were incubated for 1h with EdU in sterile 1XPBS. Liver slice or cells were fixed in PFA 4% and block/permeabilize for 30 minutes in 1xPBS 3% BSA + 0.2% Triton (1% Triton for liver slices). EdU reaction mix (100mM Tris pH 8.5, 4mM CuSO4, 625 nM Alexa Azide, 100mM Ascorbic acid) was incubated for 30 minutes.

### Histology and immunohistochemistry

Human and mouse liver specimens were fixed overnight in 4% paraformaldehyde and embedded in paraffin. Tumors arising in mice from different oncogene combinations and used for Fascin1 IHC were induced in FVB/N females (see references in the pertaining section of text). Sections were done at 5μm in thickness. Liver lesions were evaluated and classified by two board-certified pathologists and liver experts (S.R. and M.E.). For immunohistochemistry, slides were deparaffinized in xylene, rehydrated through a graded alcohol series, and rinsed in PBS. Antigen unmasking was achieved by boiling in 10 mM sodium citrate buffer (pH 6.0) for 10 min, followed by a 20-min cool down at room temperature. After a blocking step with the 5% goat serum and Avidin-Biotin blocking kit (Vector Laboratories, Burlingame, CA), human tissue slides were incubated with primary antibody overnight at 4°C. Slides were subjected to 3% hydrogen peroxide for 10 min to quench endogenous peroxidase activity and, subsequently, the biotin conjugated secondary antibody was applied at a 1:500 dilution for 30 min at room temperature. Immunoreactivity was visualized with the Vectastain Elite ABC kit (Vector Laboratories, Burlingame, CA), using Vector NovaRed (Vector Laboratories) as the chromogen. Slides were counterstained with hematoxylin.

### Luciferase assays

Cells were plated in 24-well plates and transfected with YAP/TAZ luciferase reporter 8XGTIIC-lux plasmid (50 ng/cm2) (Addgene 34615) together with CMV-lacZ (75 ng/cm2) to normalize for transfection efficiency based on CPRG (Merck) colorimetric assay. Transfected DNA content was kept equal using pKS Bluescript. Cells were harvested in luc lysis buffer (25 mM Tris pH 7.8, 2.5 mM EDTA, 10% glycerol, 1% NP-40). Luciferase activity was determined in a Tecan plate luminometer with freshly reconstituted assay reagent (0.5 mM D-Luciferin, 20 mM tricine, 1 mM (MgCO3)4Mg(OH)2, 2.7 mM MgSO4, 0.1 mM EDTA, 33 mM DTT, 0.27 mM CoA, 0.53 mM ATP). Each sample was transfected in two biological duplicates; each experiment was repeated independently with consistent results.

### RNA extraction and gene expression studies

Total RNA was isolated using commercial kits with DNAse treatment (Norgen). cDNA synthesis was carried out with M-MLV Reverse Transcriptase (Thermo) and oligo-dT primers. qPCR reactions were assembled with FastStart SYBR Green Master Mix (Roche) and run on a QuantStudio6 thermal cycler (Thermo). Gene expression levels for each biological sample were quantified as the mean between three technical replicates; *GAPDH* expression levels were used to normalize gene expression between samples. Primer sequences are provided in the following Table.

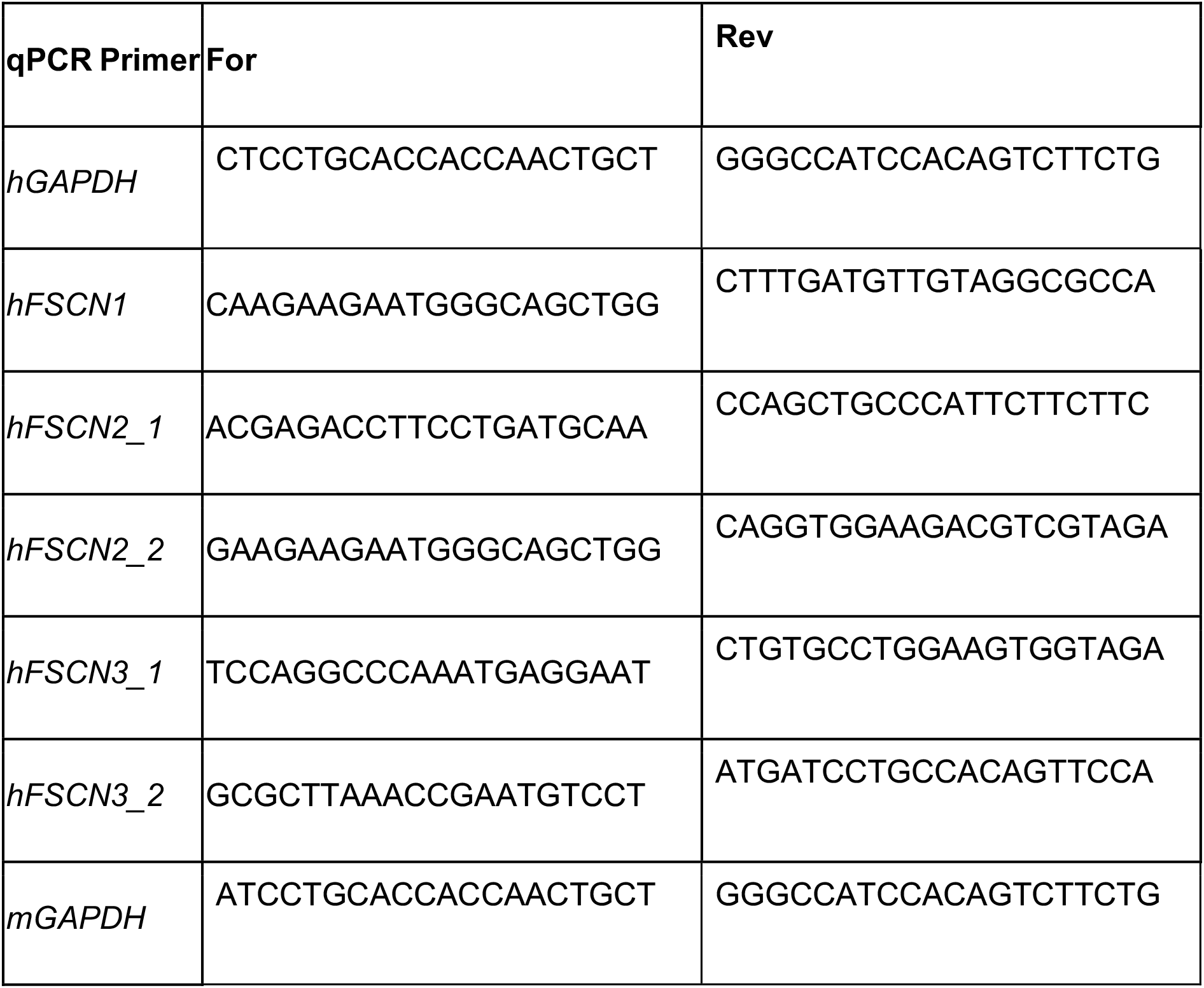
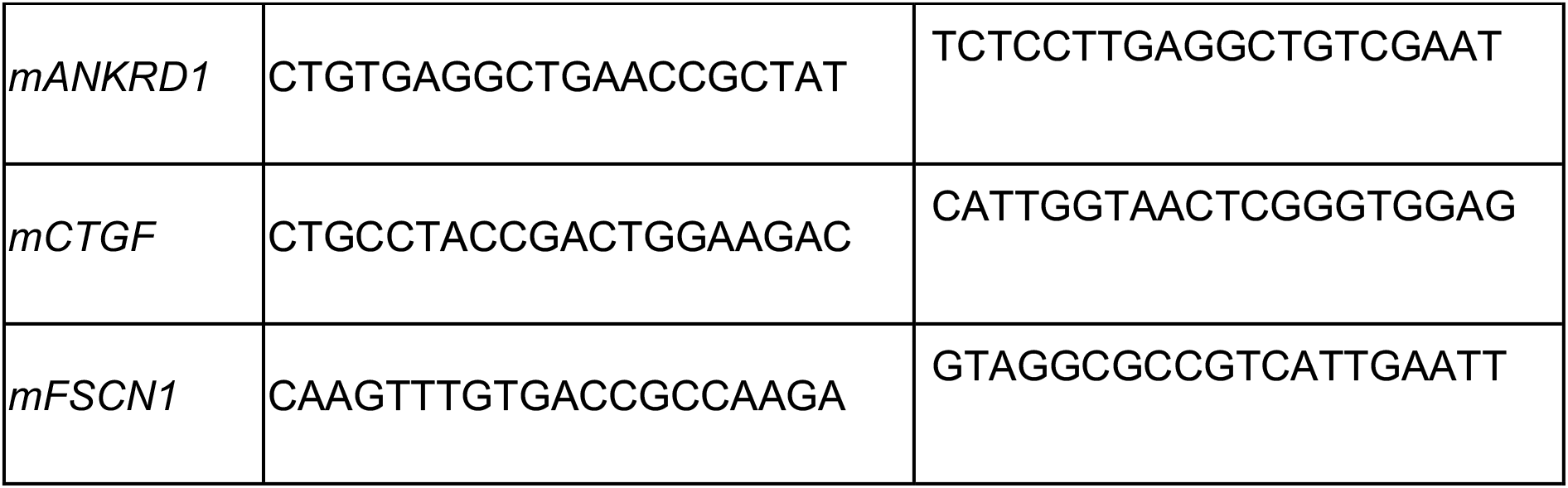

### Statistical analysis

Sample size was determined based on previous experience. Samples were not blinded for analyses. Data analyses were performed using GraphPad Prism software. Graphs indicate mean values and single points of all biological replicates (or mice), unless otherwise indicated. Data for each mouse is the average of multiple (n≥6) randomly-selected non-adjacent tissue sections. Significance was calculated by applying unpaired Welch’s t-tests between the indicated samples unless indicated otherwise in the Figure legends.

## Data availability

Data that support the findings is available in the manuscript or upon reasonable request to the corresponding author.

## Acknowledgements

We are indebted to Giorgio Scita for challenging us on the CAPZ vs. Ena/VASP antagonism, and for sharing the A4P-GFP-mito and F4P-GFP-mito plasmids; Nils Gauthier for the mCherry-Vinculin plasmid; Graziano Martello for the hyPBase plasmid; Duoja Pan for the *Yap1fl*/*fl* mice; Stefano Piccolo for the *Wwtr1(TAZ)fl/fl* mice; Antonio Rosato for help establishing the HTV technique.

---

**Supplementary Figure 1.**
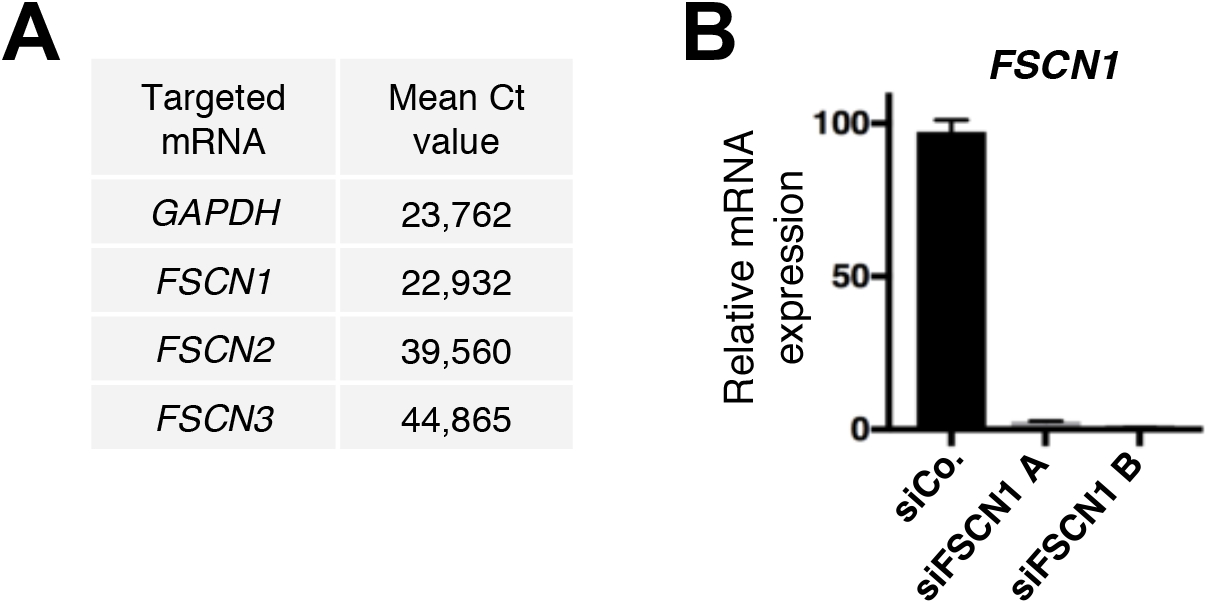
A. Mean cycle threshold (Ct) values for the indicated mRNAs in MCF10A cells, based on the same cDNA dilution. *Fascin2 (FSCN2)* and *Fascin3* (*FSCN3)* are close to the specificity detection limit. B. qPCR for *Fascin1 (FSCN1)* in MCF10A cells transfected with control siRNA (siCo.) or with two independent siRNAs targeting Fascin1 (siFascin1 A and siFascin1 B). Data are relative to *GAPDH* expression. Mean expression levels in the control sample were set to 100, and all other samples are expressed relative to this. Data are mean and s.d.

**Supplementary Figure 2.**
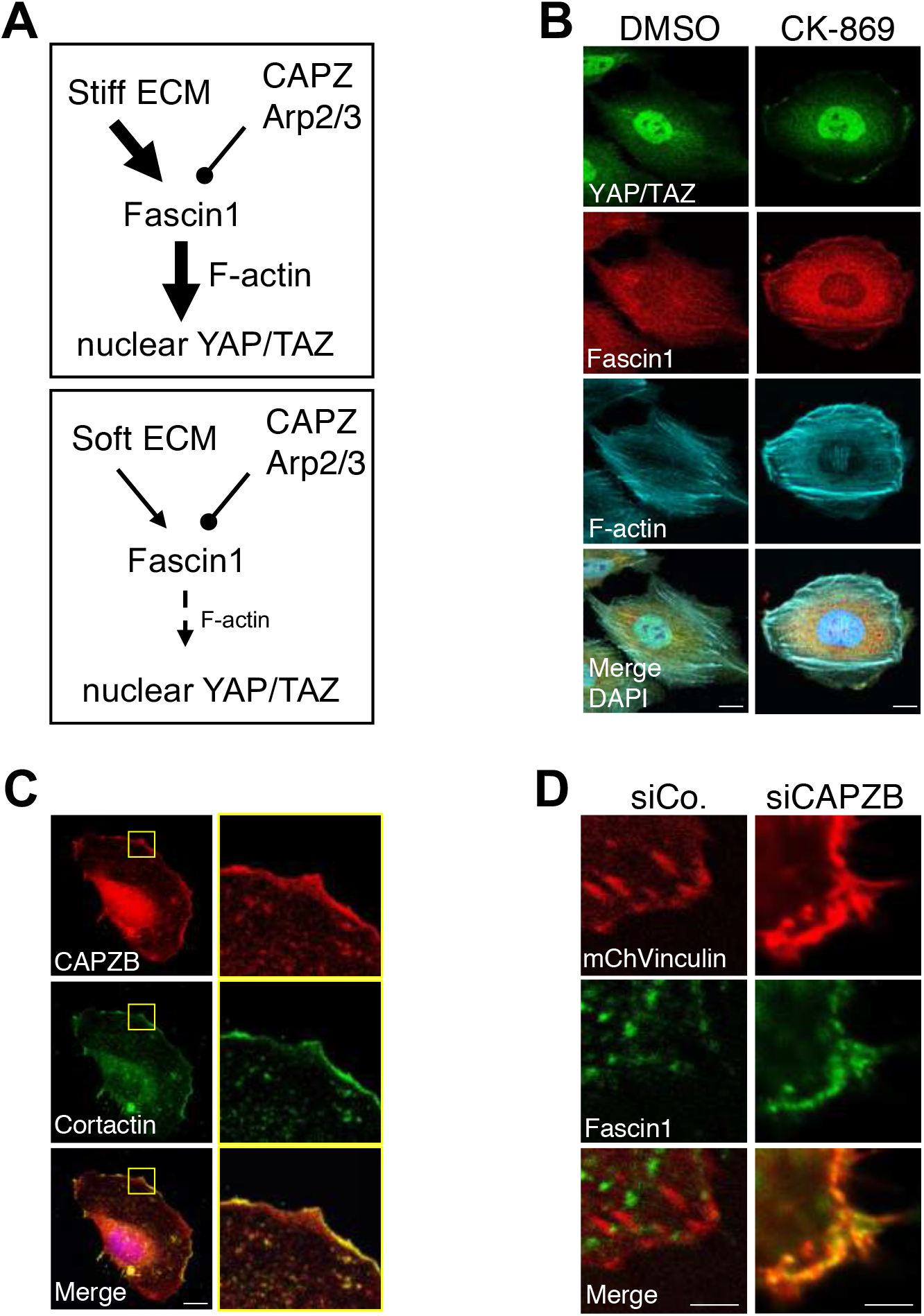
A. A simplified scheme depicting the relationships between Fascin1, ECM stiffness and CAPZ-Arp2/3. A stiff ECM sustains the formation of a contractile cytoskeleton (F-actin), which requires Fascin1; in turn this cytoskeleton promotes YAP/TAZ nuclear localization by multiple mechanisms (see introduction). In these conditions, the inhibitory effect of CAPZ and Arp2/3 are not strong enough to influence YAP/TAZ localization. On a soft ECM, decreased formation of a contractile cytoskeleton is competed by CAPZ and Arp2/3 activity, whose activity becomes relevant to regulate YAP/TAZ localization. B. Representative immunofluorescence images of MCF10A cells treated with the CK-869 Arp2/3 inhibitor and vehicle control (DMSO) and stained for Fascin1, YAP/TAZ, F-actin (phalloidin) and DAPI as nuclear counterstain. Scale bar= 10μm. C. Representative immunofluorescence images of MCF10A plated on fibronectin-coated plastics and stained for endogenous CAPZB and Cortactin, used here as a marker of cortical branched actin. Scale bar= 10μm. D. Representative immunofluorescence images of MCF10A transfected with CAPZB siRNA (siCAPZB) or control siRNA (siCo.) together with plasmid expressing mCherry-Vinculin (mChVinculin) to visualize focal adhesions (FAs). Cells were then stained for endogenous Fascin1 to evaluate co-localization with mCherry-Vinculin. Scale bar = 1μm.

**Supplementary Figure 3.**
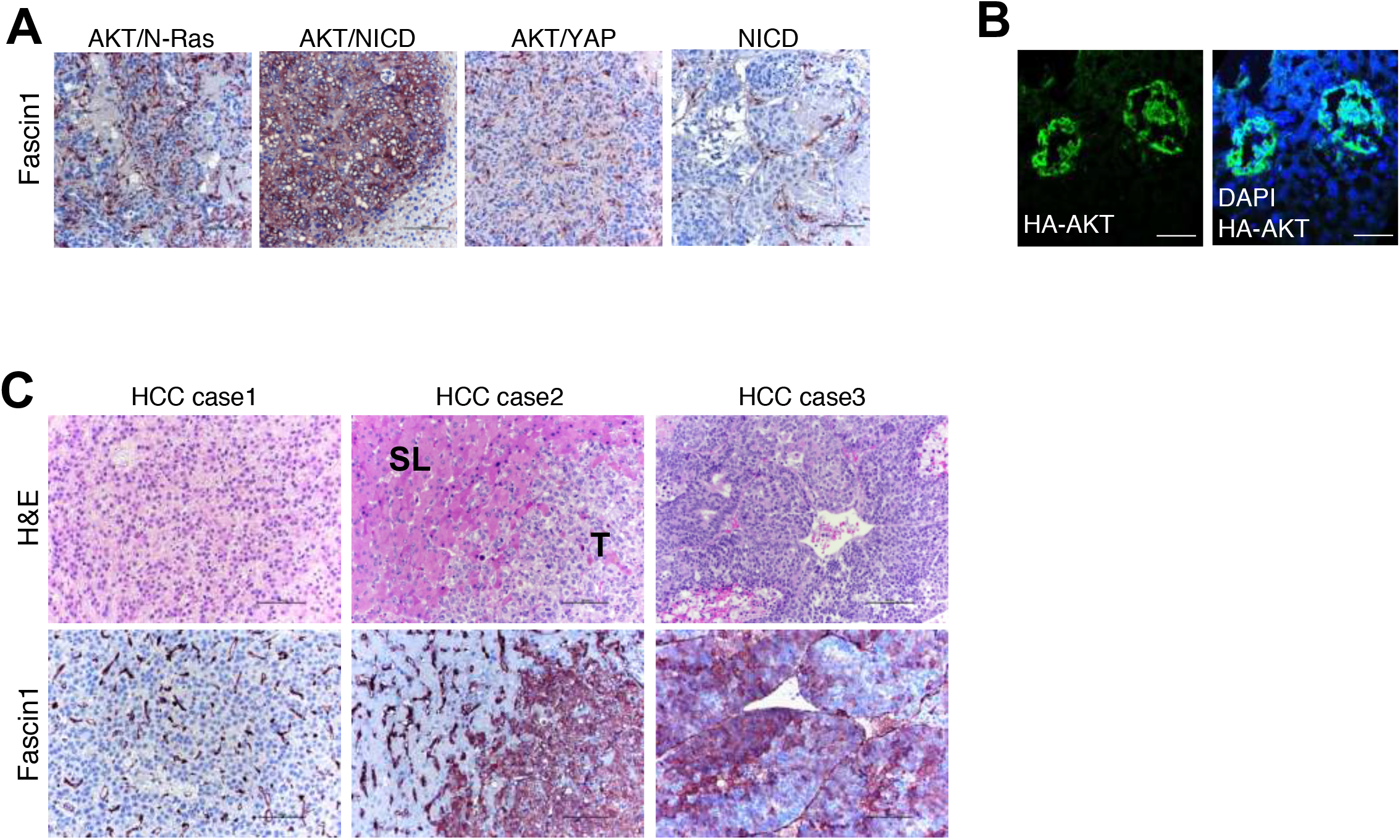
A. Higher magnifications of immunohistochemistry images shown in Fig. 4D. Original magnifications: 200x; scale bar= 100μm. B. Representative immunofluorescence images of liver sections from C57BL/6N mice transduced by hydrodynamic tail vein (HTV) injection with transposon plasmids expressing myristoylated AKT (HA-AKT), Notch Intracellular Domain (NICD) and control short-hairpin RNA (shCo.). Expression of myr-AKT together with NICD induce the formation of intrahepatic cholangiocarcinomas originating from dedifferentiated hepatocytes (Fan et al., 2012; Moya et al., 2019). Liver sections were stained for HA to localize neoplastic lesions and DAPI as nuclear counterstain. Livers were analyzed 3 weeks after HTV. Scale bar= 45 μm C. Right: a case of hepatocellular carcinoma (HCC) exhibiting immunoreactivity for Fascin1 limited to liver sinusoids and endothelial cells. Middle: a case of HCC showing intense cytoplasmic staining for Fascin1 in tumor cells (T), compared to the non-tumorous surrounding liver (SL). Left: a case of HCC displaying patchy immunolabeling for Fascin1 in tumor cells. Original magnification: 200x; scale bar= 100μm.

**Supplementary Figure 4.**
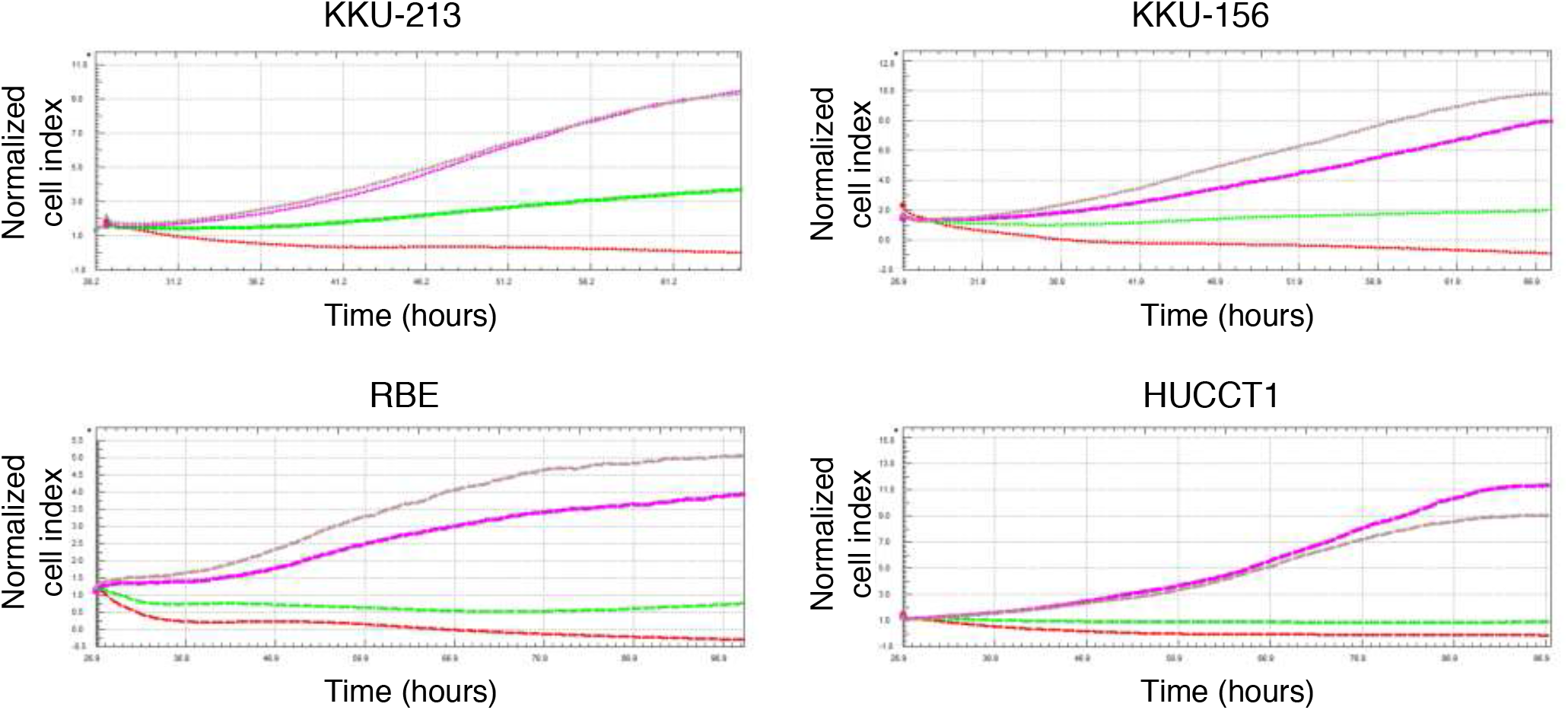
Real-time growth curve of four different human iCCA cell lines (KKU-213, KKU-156, RBE and HUCCT1) treated with different doses of G2 Fascin inhibitor, as measured by the xCELLigence Real Time Cell Analysis system (ACEA Biosciences Inc., San Diego, CA). Red curve: 100μM, green curve: 50μM, pink curve: 25μM and gray curve: 10μM. These cell lines are all positive for Fascin1 by immunoblot.

